# Darwin Assembly: fast, efficient, multi-site bespoke mutagenesis

**DOI:** 10.1101/207191

**Authors:** Christopher Cozens, Vitor B. Pinheiro

## Abstract

Engineering proteins for designer functions and biotechnological applications almost invariably requires (or at least benefits from) multiple mutations to non-contiguous residues. Several methods for multiple site-directed mutagenesis exist, but there remains a need for fast and simple methods to efficiently introduce such mutations – particularly for generating large, high quality libraries for directed evolution. Here, we present Darwin Assembly, which can deliver high quality libraries of over 10^8^ transformants, targeting multiple (> 10) distal sites with minimal wild-type contamination (lower than 0.25% of total population) and which takes a single working day from purified plasmid to library transformation. Darwin Assembly uses commercially available enzymes, can be readily automated, and offers a cost-effective route to highly complex and customizable library generation.

## INTRODUCTION

Biology, through natural enzymes, has explored and exploited a vast repertoire of chemical reactions, including many that can be harnessed for the manufacture of clinically, biotechnologically and culturally relevant molecules (1). Nevertheless, natural enzymes are optimised to their *in vivo* setting and are often unsuited for the synthesis of biological or synthetic compounds *in vitro* or in heterologous host organisms. Consequently, expression and functional optimization, or more radical engineering is often required to generate the desired enzymatic activity, whether boosting an existing activity or changing enzyme function altogether. Those needs have led to the flourishing of directed evolution and protein engineering and it has repeatedly proven possible to enhance a number of protein properties, including expression, folding(2), thermostability(3), substrate specificity(4–6) and catalytic efficiency(7, 8). While single amino acid mutations can have profound effects, they are rarely sufficient to generate the desired function and often multiple, distal mutations are required in the protein of interest – e.g. T7 RNA polymerase mutants with altered promoter recognition (R96L, K98R, E207K, E222K, N748D, P759L(9)), archaeal DNA polymerase variants capable of efficiently synthesising RNA (Y409G E664K (10)), and aminoacyl transferases used for the expansion of the genetic code (T107C, P254T, C255A (11)).

Sequential cycles of individual site mutagenesis and screening can be effective in navigating from natural to engineered catalyst (4, 5, 7, 12, 13) but such an approach requires a sequence landscape where improvement can be detected in each of the intermediates and pre-defines the evolutionary path followed(14, 15). Significantly, such iterative approaches cannot uncover phenotypes requiring epistatic mutations as multiple mutations are never made in the same round (16–18). Simultaneous multiple site mutagenesis bypasses this limitation but shifts the bottleneck to the generation of such enzyme libraries. PCR-based methods for introducing mutations to single codons, or a short patch of contiguous codons, in a gene of interest are efficient and well-established (19), but are limited to the number of mutations that can be incorporated into iPCR primers. Simultaneous introduction of multiple mutations at distal sites in a gene is significantly more challenging. A number of strategies have been developed to introduce multiple mutations into plasmid DNA (Table 1) that cover a wide spectrum in specificity (precision over which sites are targeted for mutagenesis), efficacy (fraction of mutagenized population), efficiency (number of transformants modified at all targeted positions) and complexity (number of experimental steps involved per mutagenesis cycle) – with at least one of these being inevitably compromised.

**Table 1:**
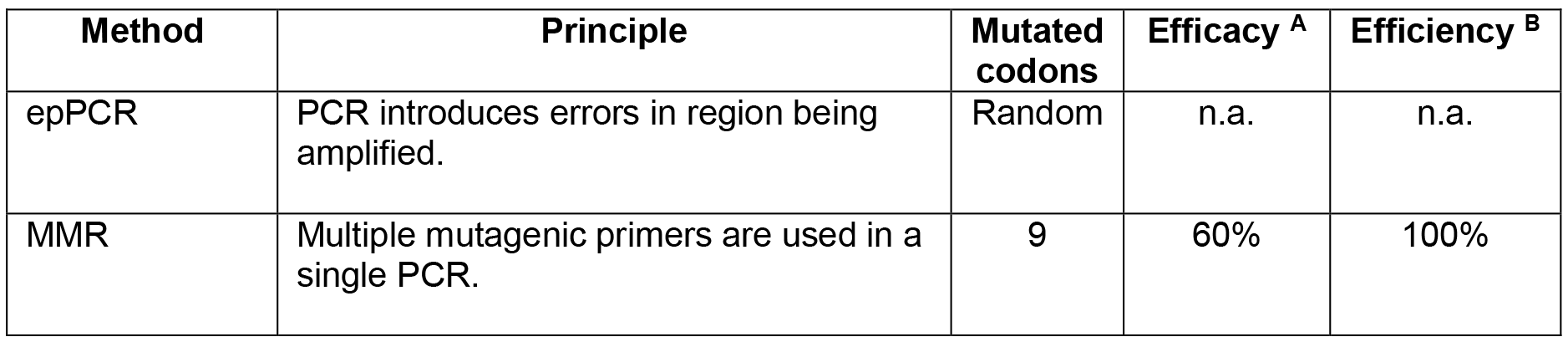
Common mutagenesis methods for the introduction of multiple distal mutations. Error-prone PCR(43) is simple but mutations introduced cannot be targeted to particular codons and its effectiveness is dependent on amplification biases and random incorporation of errors(44). In these circumstances efficacy and efficiency are not applicable concepts (n.a.). Some methods are not effective at mutating the whole population(45, 46) and/or incorporating all mutations(47, 48). Methods that can introduce mutations effectively and efficiently tend to require complex and time-consuming steps, such as phagemid propagation to generate dU-containing DNA(49, 50), chemical degradation steps(51), multiple PCR reactions followed by overlap extension PCR reactions to assemble the mutated gene(46, 52, 53) or modularization of the library into PCR-tractable libraries that can be later re-assembled via Golden Gate assembly(6). Other methods, such as TAMS(54) and OD SPM(55), are simple and effective. Both introduce diversity by using primers containing the target mutations for primer extension and ligation against single-stranded templates followed by PCR and cloning(54, 55). However, neither of these methods remove the second strand of the original plasmid and both use T4 DNA ligase, which can ligate across gaps, mispairs and cannot tolerate high temperatures favourable for specific annealing(29) – with resulting compromises in efficiency of library assembly. ^**a**^ Efficacy is defined as the fraction of the population containing mutations. ^**b**^ Efficiency is defined as the fraction of mutated clones where all targeted sites are mutated. Some methods do not report (NR) their efficacy or efficiency.

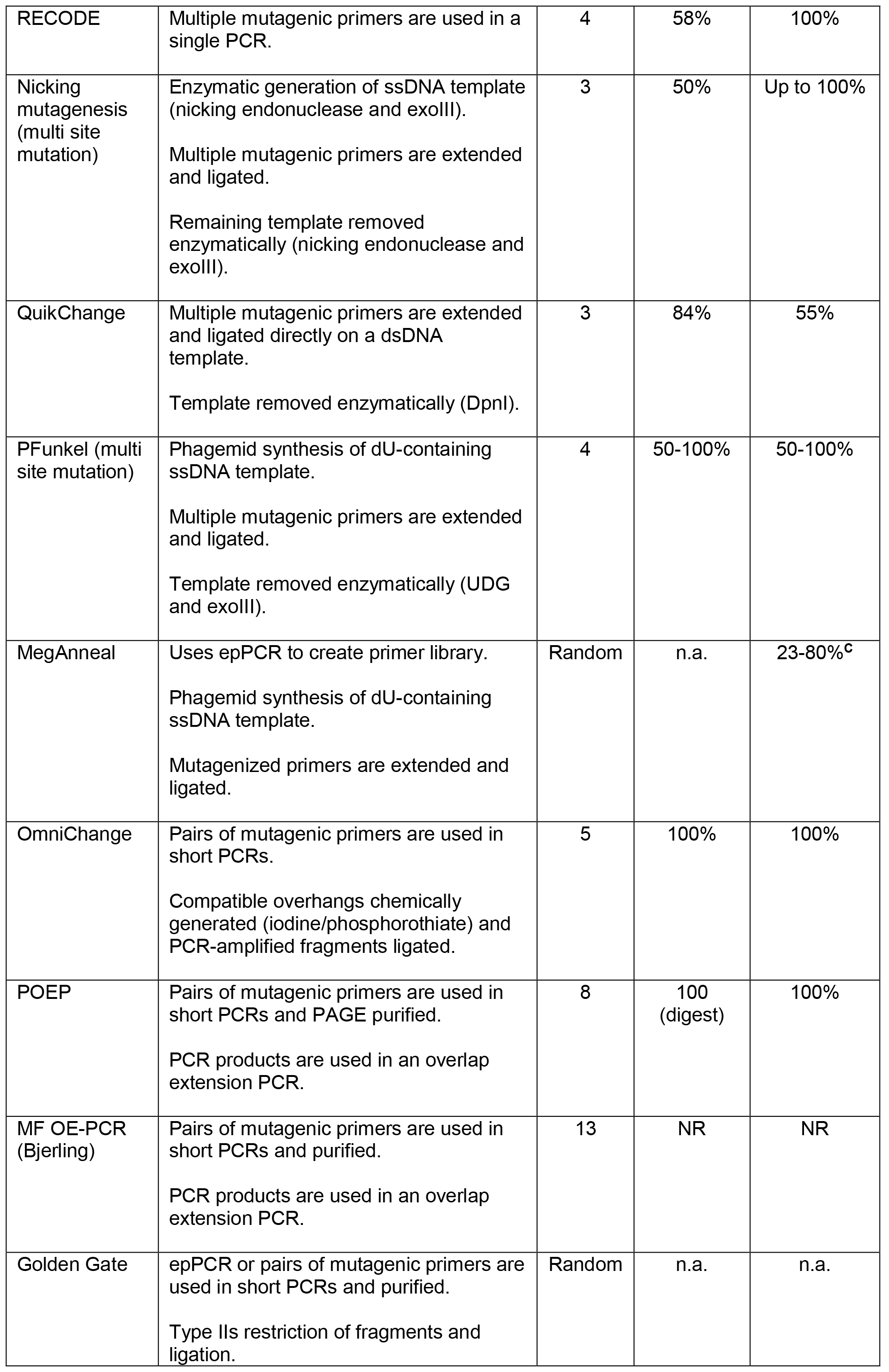

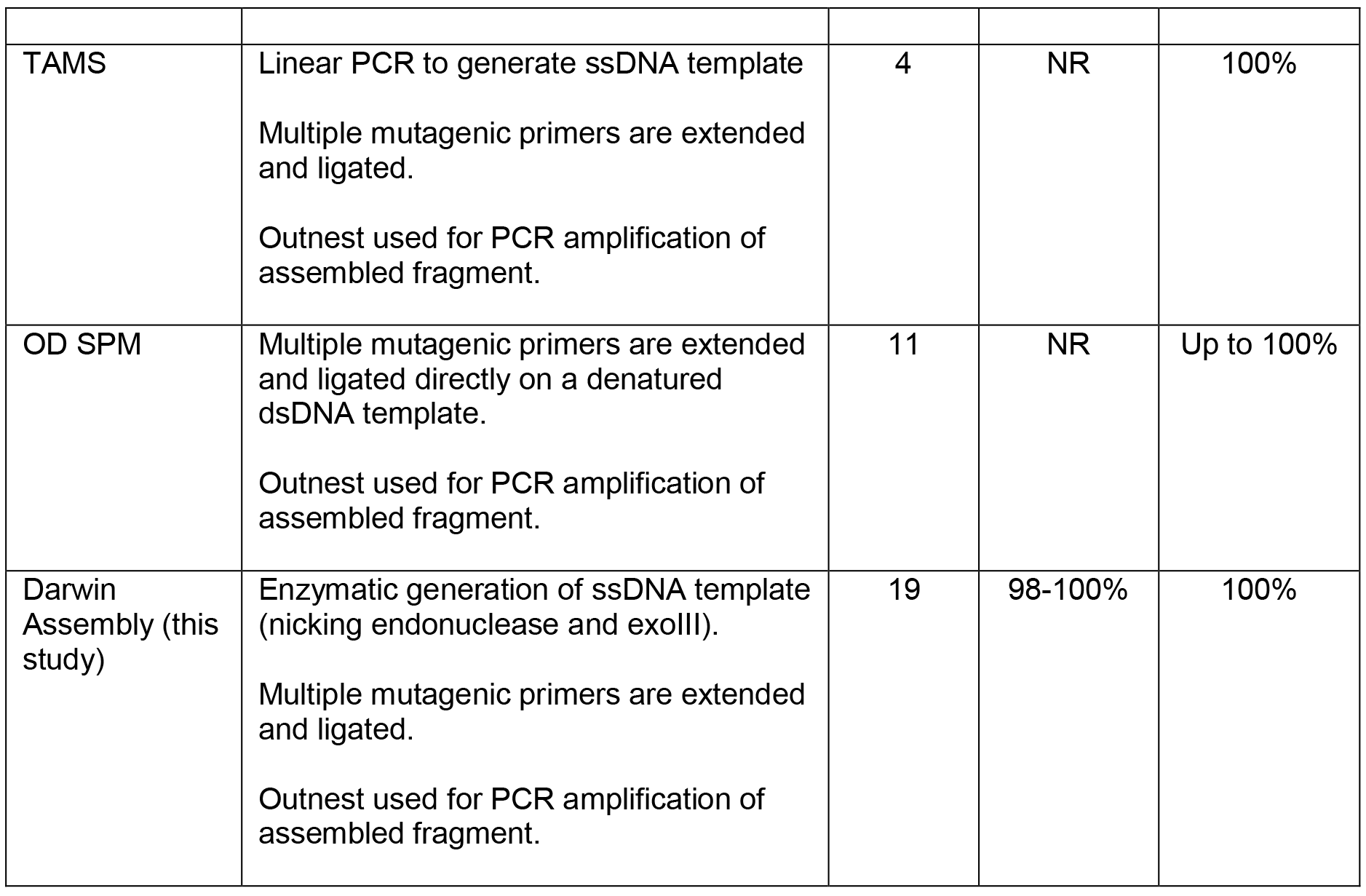

We set out to develop a new library assembly method that could uncompromisingly address all four quality bottlenecks of simultaneous multiple site saturation mutagenesis. Here, we describe Darwin Assembly – a simple, fast, low-cost and flexible platform capable of delivering large (>10^8^ transformants), user-defined libraries with any desired combination of mutations anywhere in a gene of interest, or indeed in multiple genes/features in a plasmid. Darwin Assembly uses commercially available reagents and is automation-compatible. It takes a single working day (from plasmid isolation to library transformation) and is both highly effective (less than 0.25% wild-type sequences in the generated libraries) and efficient (all clones mutated at all positions targeted).

## MATERIAL AND METHODS

### Reagents

Enzymes were from New England Biolabs (NEB; Ipswich, MA, USA) unless otherwise stated. MyTaq HS DNA Polymerase was from Bioline (London, UK). KOD Xtreme Hot Start DNA Polymerase was from EMD Millipore (Watford, UK). All oligonucleotides were from Integrated DNA Technologies (IDT; Coralville, IA, USA). ATP and NAD were from NEB. DTT was from Promega (Madison, WI, USA). PEG 8000 was from Fisher Scientific (Loughborough, UK). dNTPs were from Bioline. PCR products were purified using GeneJET PCR Purification Kits (Thermo Fisher Scientific, Waltham MA, USA), Nucleospin Gel and PCR Clean-up (Machery-Nagel GmbH, Düren, Germany) or Monarch PCR and DNA Cleanup kits (NEB). Gel purification was carried out using Monarch DNA Gel Extraction Kit (NEB). Unless otherwise stated, all site-directed mutagenesis was carried out by iPCR(19) followed by PCR purification column cleanup and blunt end ligation. All oligonucleotide sequences are provided in Table S1 and all plasmids sequences are provided as Supplementary Information.

### Plasmids and cells

Plasmid pSB1C3A2 was made from pSB1C3 (http://parts.igem.org/Part:pSB1C3) by inserting a beta-lactamase gene for ampicillin resistance (plasmid pSB1C3A) and removing an unwanted Nb.BtsI restriction site by site-directed mutagenesis, so that there were only nicking sites on one strand.

Plasmid pRST.AS11B.AS3.4(20), encoding *Saccharomyces cerevisiae* tryptophanyl tRNA synthetase, and pUC_T7RSS(21) were the kind gifts from Andrew Ellington (University of Texas at Austin, Texas, USA). pRST.AS11B.AS3.4 was mutated to remove BsaI sites (plasmid pRST.11C.AS3.4), which would interfere in the cloning post-assembly. T7RSS (encoding T7 RNA polymerase “reduced secondary structure”) was subcloned into a pBAD30(22) modified by site-directed mutagenesis to remove its BsaI sites. Site-directed mutagenesis was also carried out to remove a BspQI site within the T7RSS open reading frame by introducing a silent mutation (T7RSS gene A1671G), and to introduce 2 BspQI sites in the pBAD30 vector immediately downstream of the T7RSS gene (plasmid pBAD3-b2_T7SS in Supplementary Information).

Plasmid pET29aΔΔΔ_TgoT encoded a codon-optimised engineered archaeal B-family DNA polymerase from *Thermococcus gorgonarius*(23) (GeneWiz, UK) in a modified pET29a(+) vector lacking *rop* and harbouring a modified translation initiation sequence engineered to remove S-tag and thrombin cleavage sites. Plasmid pGDR11-KOD, encoding KOD (archaeal B-family DNA polymerase from *Thermococcus kodakarensis)*, was the kind gift from John Chaput (University of California, Irvine, USA).

*E. coli* NEB 10-beta or T7 Express lysY/I^*q*^ (NEB) were used in all experiments, with transformation using electrocompetent cells and following manufacturer’s recommendations. Plates were LB agar and liquid media was 2xTY. Antibiotics were used at 100 μg/ml (ampicillin), 33 μg/ml (chloramphenicol) or 50 μg/ml (kanamycin).

### Oligonucleotide design

The oligonucleotides used for Darwin Assembly can be divided into three groups: mutagenic primers (“inner” assembly oligonucleotides), assembly boundary oligonucleotides and outnest PCR primers – see Figure 1 for details. All oligonucleotide sequences used in this work are summarised in Table S1. “Inner” mutagenic primers were designed to have annealing temperatures between 55 and 60°C (assuming no mismatches) and to have at least 11 nucleotides to each side of a mutation (in some cases extended until a G or C was reached) to ensure efficient primer binding to the single-stranded template and efficient ligation by Taq DNA ligase.

Boundary oligonucleotides form the 5’- and 3’-ends of the strand generated during isothermal assembly (Figure S1). The boundary oligonucleotides were designed to anneal to the same strand at each end of the region being assembled and to harbour overhangs. Mutations can be introduced in the annealing sequences (same design as above for ‘inner’ mutagenic primers), but usually were not. Overhangs typically include restriction sites and outnesting PCR priming sites. Three variants of assembly boundary primers were used in the work. The first consisted to two oligonucleotides, the 5’-oligonucleotide containing no modifications and the 3’-oligonucleotide having a protected 3’-end (3’inverted-dT) to protect the single stranded 3’ overhang from exonuclease degradation during the assembly reaction (Figure S1). This method was efficient for short assemblies (< 1.0 kb) but longer assemblies (2-3 kb) required modifications.

The second variant tested was based on purification of the mutated strand generated during the assembly reaction by biotin-streptavidin pulldown. Here, two oligonucleotides were used: a 5’-oligonucleotide harbouring a biotin-TEG modification at its 5’-end and a 3’-oligonucleotide with a protected 3’-end (Figure 1). We presume that the gains in efficiency observed are due to improved purification of the assembled DNA from mutagenic primers and single-stranded plasmid template.

**Figure 1.**
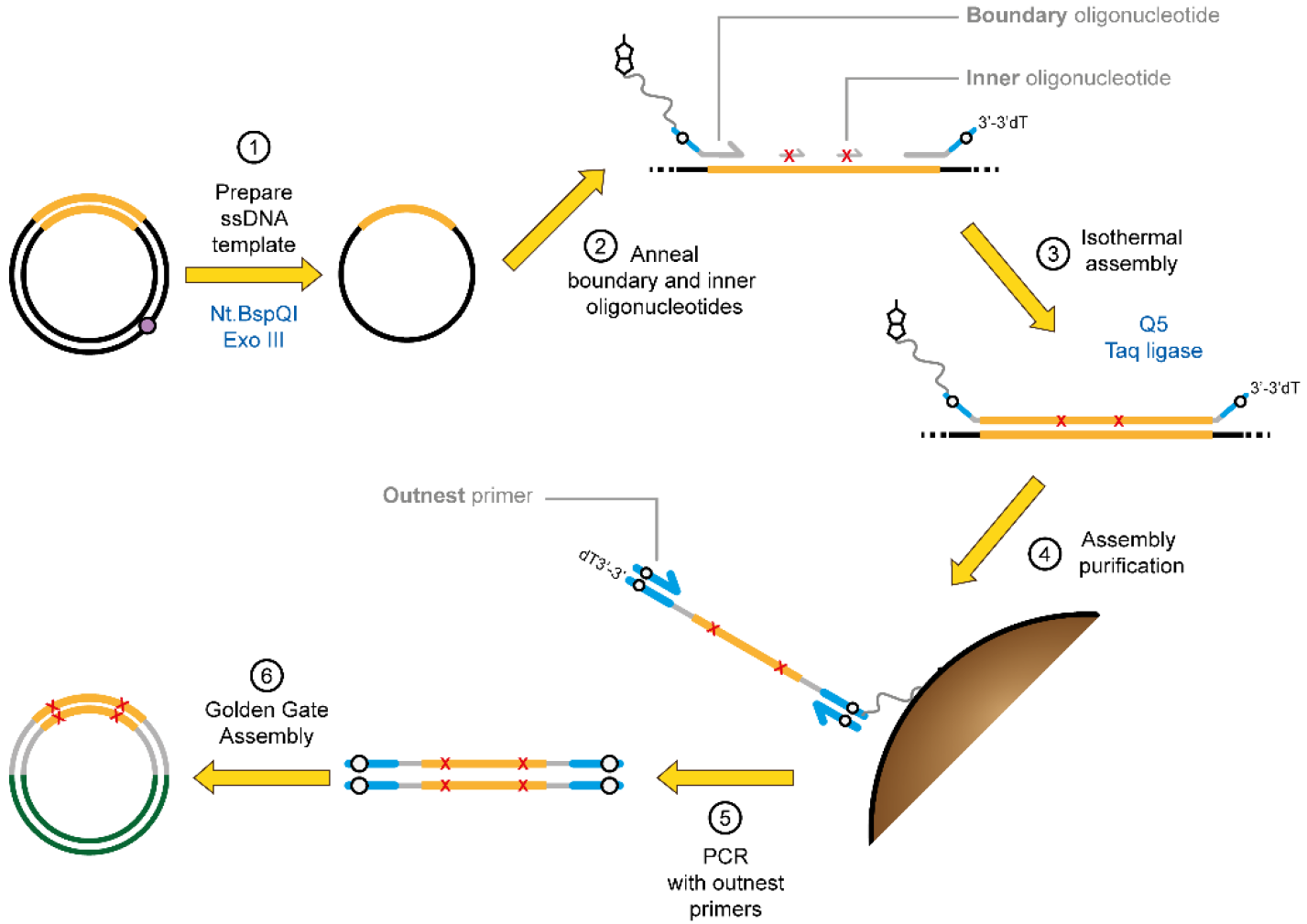
Principles of Darwin Assembly. Plasmid DNA (black, with the gene of interest in orange) is nicked by a nicking endonuclease (at the purple dot) and the cut strand degraded by exonuclease III (1). Boundary and inner (mutagenic) oligonucleotides are annealed to the ssDNA plasmid (2). Key features of the oligonucleotides are highlighted: 5’-boundary oligonucleotide is 5’-biotinylated; non-complementary overhangs are shown in blue with Type IIs endonuclease recognition sites shown in white; mutations are shown as red X in the inner oligonucleotides; 3’-boundary oligonucleotide is protected at its 3’-end. After annealing, primers are extended and ligated in an isothermal assembly reaction (3). The assembled strand can be isolated by paramagnetic streptavidin-coated beads (4) and purified by alkali washing prior to PCR using outnested priming sites (5) and cloning (6) using the type IIS restriction sites (white dots). The purification step (4) is not always necessary but we found it improved PCR performance, especially for long assembly reactions (>1 kb).

The third variant tested used a single, long oligonucleotide harbouring both 5’ and 3’ binding sites, akin to a padlock probe(24), with sites for post-assembly amplifications (i.e. restriction sites and outnest PCR priming sites) linked by a short flexible linker (dT_5_). This topology, referred to as the θ oligonucleotide, generates a closed circle after isothermal assembly, allowing exonuclease cleanup and hence removal of partial assembly products, unincorporated oligonucleotides and original plasmid (Figure 2).

**Figure 2.**
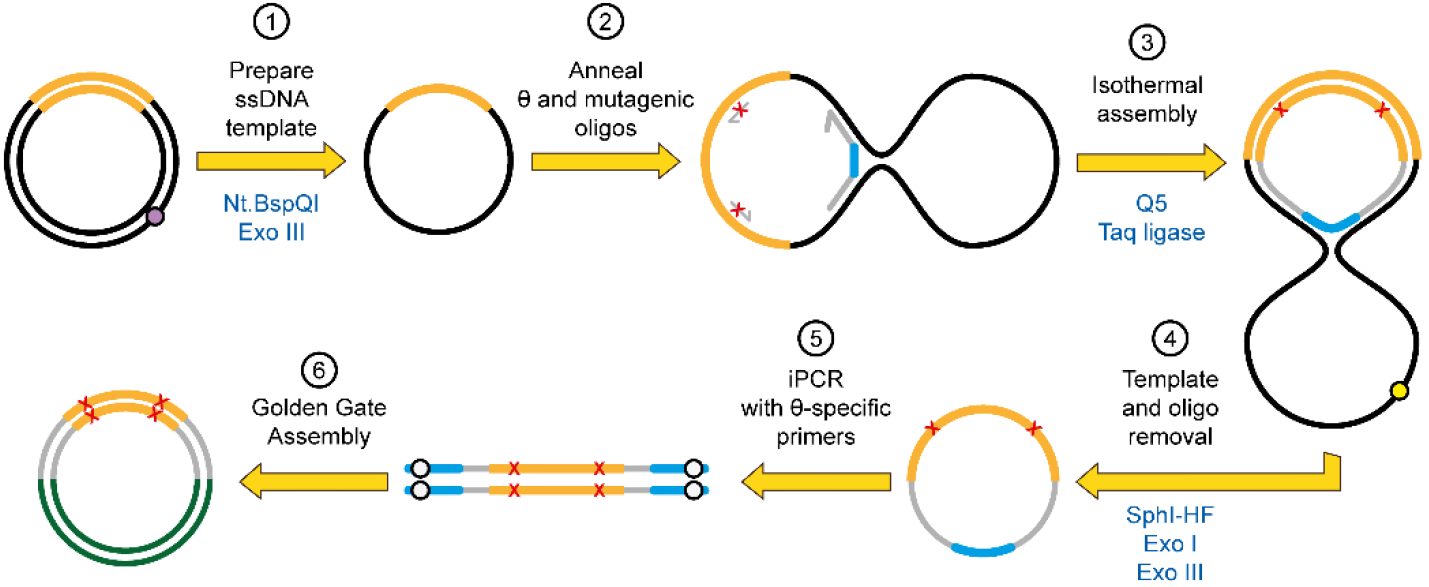
Darwin Assembly using a θ oligonucleotide. Here, a single θ oligonucleotide is used in place of the two boundary oligonucleotides allowing enzymatic cleanup after the assembly reaction. Plasmid DNA (black, with the gene of interest in orange) is nicked by a nicking endonuclease (at the purple dot) and the nicked strand degraded by exonuclease III (1). Inner oligonucleotides and a single θ oligonucleotide are annealed to the ssDNA plasmid (2). The θ oligonucleotide encodes both assembly priming and termination sequences linked by a flexible linker such that successful assembly of the mutated strand results in a closed circle (3). The template plasmid can now be linearized (e.g. at the yellow dot, by adding a targeting oligonucleotide and appropriate restriction endonuclease) and both exonuclease I and exonuclease III added to degrade any non-circular DNA (4). The mutated gene can now be amplified from the closed circle by PCR (5) and cloned into a fresh vector (6) using the type IIS restriction sites (white dots).

Outnest PCR primers, designed to anneal onto the assembly boundary primer overhangs, were typical PCR primers used to amplify the synthesised libraries prior to cloning by Golden Gate assembly(25).

### Single strand plasmid generation

Plasmid DNA, isolated from overnight bacterial cultures using GeneJET Plasmid Miniprep Kits (Thermo Fisher Scientific), was made single stranded by co-incubation with a nicking endonuclease and exonuclease III (exoIII). Typically, DNA was digested at 50-60 nM (90 to 235 ng/μl, depending on plasmid size) with 3 to 10 U of nicking enzyme and 40 to 60 U of exoIII per μg of DNA. Plasmid DNA was vacuum concentrated (SpeedVac, ThermoFisher Scientific) if necessary prior to digestion. Reactions, in 1x CutSmart buffer (NEB), were carried out for 2 h at 37°C, followed by 20 min at 80°C to inactivate the enzymes. Nt.BspQI and Nb.BtsI were used successfully but Nt.BbvCI was consistently less active under the conditions used. No other nicking endonucleases have been tested. Digestion was confirmed by agarose gel electrophoresis followed by SYBR Gold (Thermo Fisher Scientific) staining. ssDNA was used for assembly without further purification and was not re-quantified.

### Oligonucleotide phosphorylation

Mutagenic primers and the 3’ boundary assembly oligonucleotides were phosphorylated at their 5’-ends using T4 polynucleotide kinase (T4 PNK, NEB), to enable ligation during isothermal assembly. Oligonucleotides, at 4 – 20 μM final oligonucleotide concentration, were phosphorylated in 1x CutSmart buffer supplemented with 1 mM ATP and 0.1 U/μl T4 PNK. Typically, the inner mutagenic oligonucleotides were mixed and phosphorylated in a single reaction and the 3’ boundary oligonucleotide phosphorylated separately. Reactions were carried out for 2 h at 37°C, followed by 20 min at 80°C, to inactivate the enzymes. Phosphorylated oligonucleotides were added to subsequent assembly without further purification.

### Isothermal Assembly

Firstly, single-stranded plasmid DNA (typically 0.1 to 0.2 pmol per reaction) was mixed with mutagenic oligonucleotides (at 100 to 250-fold molar excess) and boundary assembly oligonucleotides (at 10 to 50-fold molar excess over plasmid DNA). Plasmid and oligonucleotide mixtures, typically between 3 and 5 μl, were then annealed by freezing samples at −20°C for at least 15 min and subsequently returning to room temperature. Alternatively, efficient annealing was also obtained by heating samples for 5 min to 95°C and cooling them at 0.1°C/s to 4°C.

After annealing, 1 volume 2x Darwin Assembly enzyme mix (2x DA mix) was added and the reaction incubated at 50°C for 10 min to 1 h, depending on the length of the assembly (60°C was also successfully tested, data not shown). Ten minutes is sufficient even for long assemblies (e.g. T7 RNA polymerase; 2.7 kb) although slightly higher PCR yields are seen after longer reaction times. 2x DA mix consists of: 0.05 U/μl Q5 High-Fidelity DNA polymerase (not hot start), 8 U/μl Taq DNA ligase, 2 mM NAD+, 0.4 mM each dNTP, 10% (w/v) PEG 8000, 2 mM DTT and 1x CutSmart buffer. This reaction mix is closely modelled on the Gibson Assembly (26), with Phusion replaced by Q5 DNA polymerase and the reaction buffer replaced with CutSmart buffer. It is stored frozen (and like commercially available Gibson Assembly mixes, will freeze solid).

### Assembly purification with streptavidin-coated paramagnetic beads

Streptavidin-coated paramagnetic beads were used to clean up assemblies carried out using 5’-biotinylated 5’-boundary oligonucleotides.

Initially, 5 μl Dynabeads MyOne Streptavidin C1 beads (Thermo Fisher Scientific) were blocked in 2x BWBS-T (20 mM TRIS·HCl pH 7.4, 2 M NaCl, 0.2% v/v Tween-20, 2 mM EDTA) for over an hour on a spinning wheel rotator at room temperature. The beads were isolated on a magnetic stand and resuspended in 50 μl 2x BWBS-T (20 mM TRIS·HCl pH 7.4, 2 M NaCl, 2 mM EDTA, 0.2 (w/v) Tween-20).

Once the assembly reaction was completed, its volume was adjusted to 50 μl with PCR-grade water and transferred to a 1.5 ml tube. The pre-blocked beads were added to the reaction and mixed thoroughly. Assembly binding to beads was carried out for 3 h at room temperature, although it was later found that 10 minutes were sufficient (albeit at slight cost to yield). After capture, beads were isolated on a magnetic stand and washed twice in 200 μl 37°C 30 mM NaOH and once in 200 μl EB-T (10 mM TRIS.HCl pH 8.8, 0.1 mM EDTA, 0.01% Tween-20). Alkali denaturation is carried out at 37°C to improve bead handling – this is not essential for plasmid denaturation. The wash with EB-T is included to neutralise the beads and to reintroduce Tween-20, to prevent the beads adhering to tube walls or aggregating. After washing, the beads were resuspended in 10 μl EB (10 mM TRIS HCl pH 8.8) and used directly for PCR.

### Assembly purification by exonuclease

Exonuclease cleanup was used in conjunction with the θ oligonucleotide method for one-pot purification of assembly products. After assembly, a master mix consisting of exonuclease I (exoI), exoIII, a targeting oligonucleotide (see below) and an appropriate restriction endonuclease is added to the assembly reaction. The targeting oligonucleotide is designed to anneal efficiently to a suitable endonuclease restriction site present in the ssDNA template (e.g. SphI in pET29), so that the chosen endonuclease can cleave the local dsDNA template generated. This creates free 3’ and 5’-ends enabling exoIII and exoI to degrade the starting template, along with any partial assembly products and free oligonucleotides.

For the TgoT experiment, a 5x master mix consisting 4 U/μl exoI, 2 U/μl exoIII, 1 U/μl SphI-HF and 50 μM oligonucleotide pET_SphIcut was prepared, of which 4 μl were added to a 16 μl isothermal assembly reaction, giving final concentrations of 0.8 U/μl exoI, 0.4 U/μl exoIII, 0.2 U/μl SphI-HF and 10 μM pET_SphIcut. Reactions were carried out for 40 min at 37°C, followed by incubation for 20 min at 80°C to inactivate endo- and exonucleases.

### PCR amplification of assembled DNA

PCR amplification was carried out directly from the assembly reactions, or after its purification (whether exonuclease or beads). When carried out directed from assembly reactions, typically 1 μl assembly was used as template per 25 μl PCR. Primers outnest_1 and outnest_2, targeting sites on the boundary assembly oligonucleotides, were used to specifically amplify the assembled libraries (as their sequences were chosen to not be present in the original plasmid DNA template).

Multiple enzymes [MyTaq (Bioline), Q5 Hot Start DNA polymerase (NEB) and KOD Xtreme (EMD Millipore)] were successfully used for post-assembly PCR. In general, KOD Xtreme was found to be the most robust regarding successful PCR amplification. Reactions consisted 1x KOD Xtreme buffer, 0.3 μM outnest_1, 0.3 μM outnest_2, 0.4 mM each dNTP and 0.01 U/μl KOD Xtreme DNA polymerase. The cycling parameters used were 2 min 95°C, followed by 20-28 cycles of 15 s at 98°C, 15 s at 64°C and 2 min 30 s at 68°C, with a final polishing step of 2 min at 72°C. These conditions were typical for 2-3 kb PCR products, and extension time was adjusted as per manufacturer recommendations for shorter assemblies.

After 25 μl pilot PCRs, PCRs were scaled up (usually to 2 × 50 μl reactions) and gel purified using Monarch PCR and DNA cleanup kits (NEB). These were preferred as they allow low elution volumes and result in clean DNA (low A230 contamination). The purified products were digested with BsaI-HF to remove outnesting priming sites and generate suitable overhangs for cloning, and again purified to remove enzyme, salts and cleaved primers. DNA concentration was determined by spectrophotometry (SPECTROstar Nano, BMG Labtech, UK).

### Vector amplification and cloning

Vectors were made by iPCR using Q5 Hotstart High Fidelity DNA Polymerase. Reactions consisted of 1x Q5 buffer, 1x Combinatorial Enhancer Solution (CES)(27), 0.2 mM each dNTP, 0.5 μM each primer and 0.01 U/μl Q5 HS DNA polymerase. The PCR conditions used were 1 min at 98°C, followed by 30 cycles of 15 s at 98°C, 15 s at 60°C and 3 min at 72°C. A final polishing step extension of 1 min at 72°C was routinely included. Annealing temperature and extension time varied according to the primer pair and the size of vector being prepared.

Amplification reactions were purified, and treated concomitantly with BsaI-HF, rSAP and DpnI to create the compatible overhangs, dephosphorylate the vector and degrade any contaminating original template. The amplified vector DNA was then purified using PCR purification columns (Nucleospin, Machery Nagel) and quantified as above. For point mutations, 20 μl ligations containing 50 fmol insert, 12.5 fmol vector and 0.5 U/μl T4 DNA ligase were ligated for 10 min to 2 h at room temperature in 1x T4 DNA ligase buffer. For libraries, typically 300 fmol insert was ligated into 75 fmol vector at room temperature for 2 h in 100 μl reactions, using 0.8 U/μl T4 DNA ligase and supplemented with 0.5 U/μl of 5’deadenylase to maximise ligation efficiency.

All ligations were phenol:chloroform:isoamyl alcohol purified, isopropanol precipitated, resuspended in 5 to 10 μl EB and transformed into electrocompetent NEB 10beta (CAT, KOD and T7RSS) or NEB T7 Express LysY/I^*q*^ (TgoT) *E.coli*.

### Illumina sequencing and data analysis

All deep sequencing was carried out on an Illumina MiSeq at the UCL Genomics Centre using a 150 cycle MiSeq Reagent Kit v3 (Illumina). Libraries were prepared by PCR using KOD Xtreme Hot Start DNA Polymerase (EMD Millipore) to append barcodes and sequencing primers. Libraries were gel purified on agarose gels stained with SYBR safe (Life Technologies), visualised using a blue light source and purified using Monarch DNA Gel Extraction kits (NEB). Libraries were then quantified using a Qubit Fluorometer (Thermo Fisher Scientifc).

Codon frequencies were counted in R version 3.3.0 (28) using RStudio version 1.0.136 for Mac OS X after data processing. Raw FASTQ files were obtained from the Miseq and quality-filtered, trimmed, barcode-split and converted to FASTA using the FASTX-Toolkit (http://hannonlab.cshl.edu/fastx_toolkit/index.html). If necessary, Prinseq (http://prinseq.souceforge.net/) or TAGCleaner (http://tagcleaner.sourceforge.net/) were used to omit reads below a minimum or above a maximum length. FASTA headers were removed, reading frames aligned and the DNA sequence split into triplet codons to facilitate reading into R. More detail on the commands used is available in the Supplementary Methods.

## RESULTS

### Principles of Darwin Assembly – fast, efficient, defined multi-site assembly

The Darwin Assembly process can be split into three steps: the generation of single-stranded DNA template, isothermal assembly, and subsequent amplification for cloning (Figures 1 and 2). Each step was designed such that the output of one step becomes the input of the subsequent step without the need for purification – minimising sample loss and handling time. The result is a fast protocol that goes from plasmid DNA to library within a day and it is compatible with automation.

Single-stranded template is generated from purified plasmid DNA by the coupled reaction of a nicking endonuclease and exonuclease III (exoIII). The endonuclease specifically cuts one strand of plasmid DNA and exoIII selectively and efficiently degrades the nicked DNA strand, leaving behind a circular single-stranded template for assembly. Nevertheless, we found it increased the robustness of the assemblies to position a nicking site shortly after the 3’-end of the assembly reaction in some instances (data not shown). Nicking endonuclease and exoIII are heat inactivated and the single-stranded template can be used directly in assembly, as all enzymes used have activity in a common buffer.

For assembly, primers (except for the 5’ boundary assembly oligonucleotide) are phosphorylated enzymatically and added to the assembly reaction in the desired molar ratios. Mutations are introduced by the oligonucleotides and primer excess ensures efficient binding to ssDNA template, contributing to the high efficiency of the Darwin Assembly. Boundary assembly primers are used at lower concentrations than mutagenic primers (but still in excess of ssDNA template) to limit side reactions that could undermine a successful assembly. Primer concentration was not extensively optimised as the concentrations used proved robust for assembly and generated minimal background or unwanted assemblies.

Once oligonucleotides and ssDNA template are annealed, assembly is carried out with a non-strand displacing thermostable DNA polymerase (Q5) and a thermostable DNA ligase (Taq DNA ligase), akin to Gibson assembly(26). The high reaction temperature and the choice of DNA ligase both contribute to the specificity of the assembly, limiting off-target annealing of the primers and limiting ligation to nicks on the nascent DNA strand(29), respectively.

Depending on the complexity of the assembly, no purification may be necessary, with assembly reactions used directly as PCR templates. Purification, whether based on nucleases or isolation of biotinylated DNA post-assembly, improves the quality of subsequent PCR amplification making it less likely to contain secondary amplification products (data not shown). We assume the main source of secondary PCR products is carryover of inner assembly oligonucleotides to the PCR, where they act as primers. Reducing the concentration used in the assembly reaction may remove the need for purification but at the possible risk of missing out some mutations.

PCR amplification post-assembly generates sufficient material for Golden Gate cloning(30), ensuring highly efficient and scarless cloning of the assembled fragment. Our deep sequencing results show that 98 - 100% of targeted codons are mutated as expected. No wild-type clones were detected by Sanger sequencing, even in the absence of any purification steps, and deep sequencing of biotin-purified assembled libraries showed wild-type contamination was always below 0.25%, a level comparable with sequencing error rates.

In developing the methodology and to explore its potential applications, we have applied Darwin Assembly to different genes, including chloramphenicol acetyl transferase (CAT), *Saccharomyces cerevisiae* tryptohanyl tRNA synthetase (ScWRS), *Thermococcus gorgonarius* DNA polymerase (TgoT), *Thermococcus kodakaraenis* DNA polymerase (KOD) and T7 RNA polymerase (T7RSS), which differ in length as well as composition. We demonstrate Darwin Assembly can be used to wholesale reassign codons in a gene of interest, create amino acid scanning and insertion/deletion (indel) libraries, as well as complex mutational libraries – where diversity can be highly customisable (e.g. non-redundant equimolar incorporation of triplet codons).

### Whole gene codon reassignment

We chose chloramphenicol acetyl transferase (CAT) as a suitable initial target to develop and optimise Darwin Assembly, as it is a small (660 bp), well-characterised gene. Our initial proof of principle was to reassign all leucine codons in CAT to CTG. Six oligonucleotides were used to introduce the required 10 point mutations (in seven different codons spread over 512 bases, Figure 3a). Assemblies were carried out using two boundary oligonucleotides and without any pre-amplification purification. Two assembly conditions were tested, with either 125x inner and 25x boundary oligonucleotide molar excess over plasmid, or 250x inner and 50x boundary oligonucleotide molar excess over plasmid.

**Figure 3.**
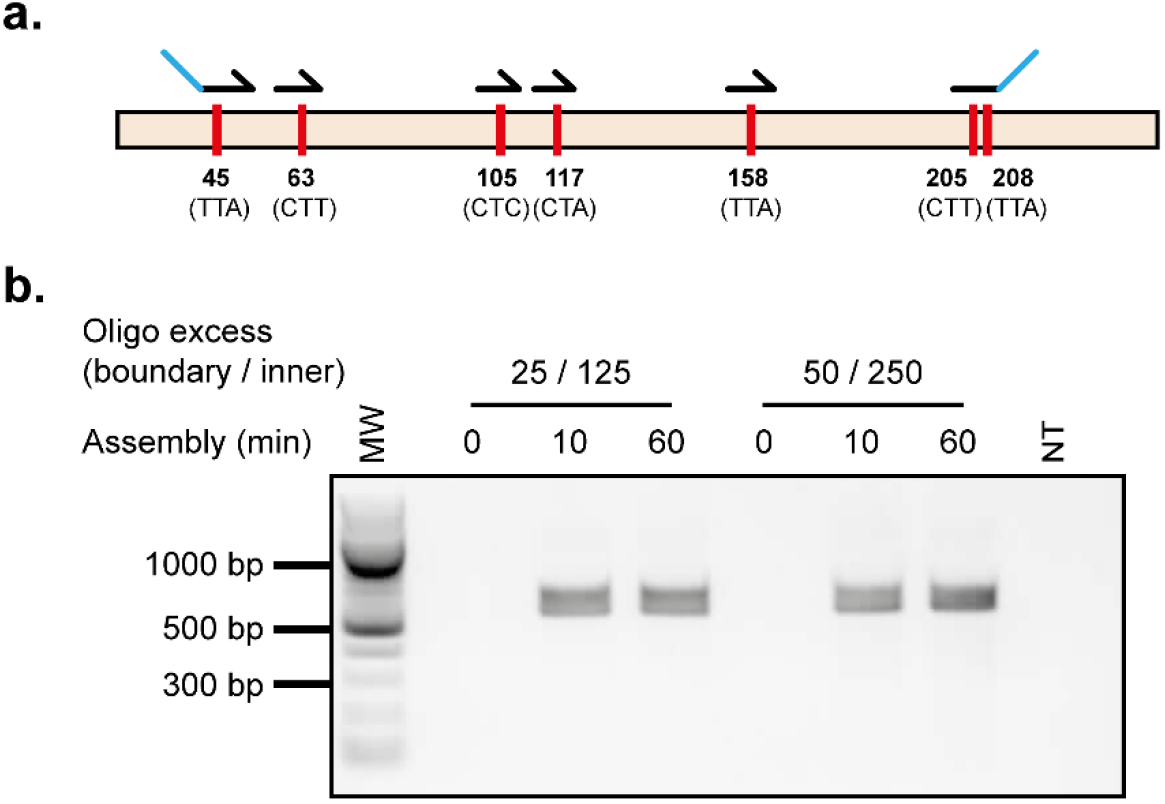
Whole gene codon reassignment. Codons targeted for reassignment and oligonucleotides used in Darwin assembly (a). Amino acid numbers are original codon sequences are shown. Outnest PCR showing amplification of the mutated CAT gene following the Darwin Assembly reaction across a range of assembly times (0, 10 and 60 minutes) and assembly oligonucleotide concentrations (b). Concentrations shown represent the molar excess of boundary (e.g. 25) and inner oligonucleotides (e.g. 125) over plasmid concentration used. Expected assembly amplicon is 568 bp. MW: 100 bp ladder (NEB).

Both assemblies were successful (Figure 3b) and 100% of the sequenced transformants (7 colonies three transformants for the lower and four of the higher concentrations) had all of the desired mutations with no mistargeting observed. In a more rigorous test of the technology, we successfully introduced 38 point mutations (in 19 different codons spanning 1963 bases) in KOD DNA polymerase in a single Darwin Assembly reaction (data not shown), using the refinements described in Materials and Methods (shown in Figure 1) to remove the initial ssDNA template and unincorporated oligonucleotides.

Encouraged by the high efficiency of the initial assembly reactions, we decided to investigate if the method was sufficiently robust and efficient for the generation of libraries of increasing complexity: site saturation (all possible mutations at a single position), partial saturation (mutually exclusive mutations targeting different sites), and indel libraries (deleting or inserting whole codons).

We tested the introduction of diversity by making a small library in *Saccharomyces cerevisiae* tryptophanyl tRNA synthetase, targeting positions important in substrate specificity (Thr107, Pro254 and Cys255(11)) with limited degeneracy (KKC, KKC and RYA, respectively). Assembly was successful and a small number of transformants were sequenced. All three sites were successfully targeted in the five sequenced transformants and no off-site mutations were observed (data not shown).

### Alanine scanning library generation

Scanning libraries introduce a single point mutation at different sites in a target gene. Alanine scanning is a traditional approach to identify functionally important residues in enzymes(31) but scanning libraries can also be used to map the local functional neighbourhood of an enzyme(32) or even to generate datasets for deep mutational scanning-guided rational protein design(33). Mutant generation can be laborious when mutations are introduced individually by site-directed mutagenesis.

We reasoned that Darwin Assembly would efficiently generate scanning libraries through the combination of oligonucleotides: all designed for the same binding site but each introducing a different mutation (Figure S2). Using the CAT gene as our model, we assembled an alanine scan library around residues Thr101 and Ser104, using Leu105 (CTC→CTG) recoding as an assembly control. Five inner oligonucleotides were designed: four introducing an alanine mutation (Thr101Ala, Phe102Ala, Ser103Ala or Ser104Ala) and one wild-type. All five oligonucleotides introduced the CTC→CTG control mutation at the Leu105 codon.

The five oligonucleotides were mixed in a 1:1:1:1:1 ratio in the assembly reaction to create a library where each variant was expected to be 20% of the final population. Assembly was successful and over 10^3^ transformants were isolated. Transformants were pooled, their plasmid DNA purified and deep sequenced - generating nearly 80,000 reads. The control Leu105 (CTC→CTG) mutation was present in 99.7% of the samples, a frequency comparable to sites that had not been targeted for mutagenesis (see Table 2).

**Table 2:**
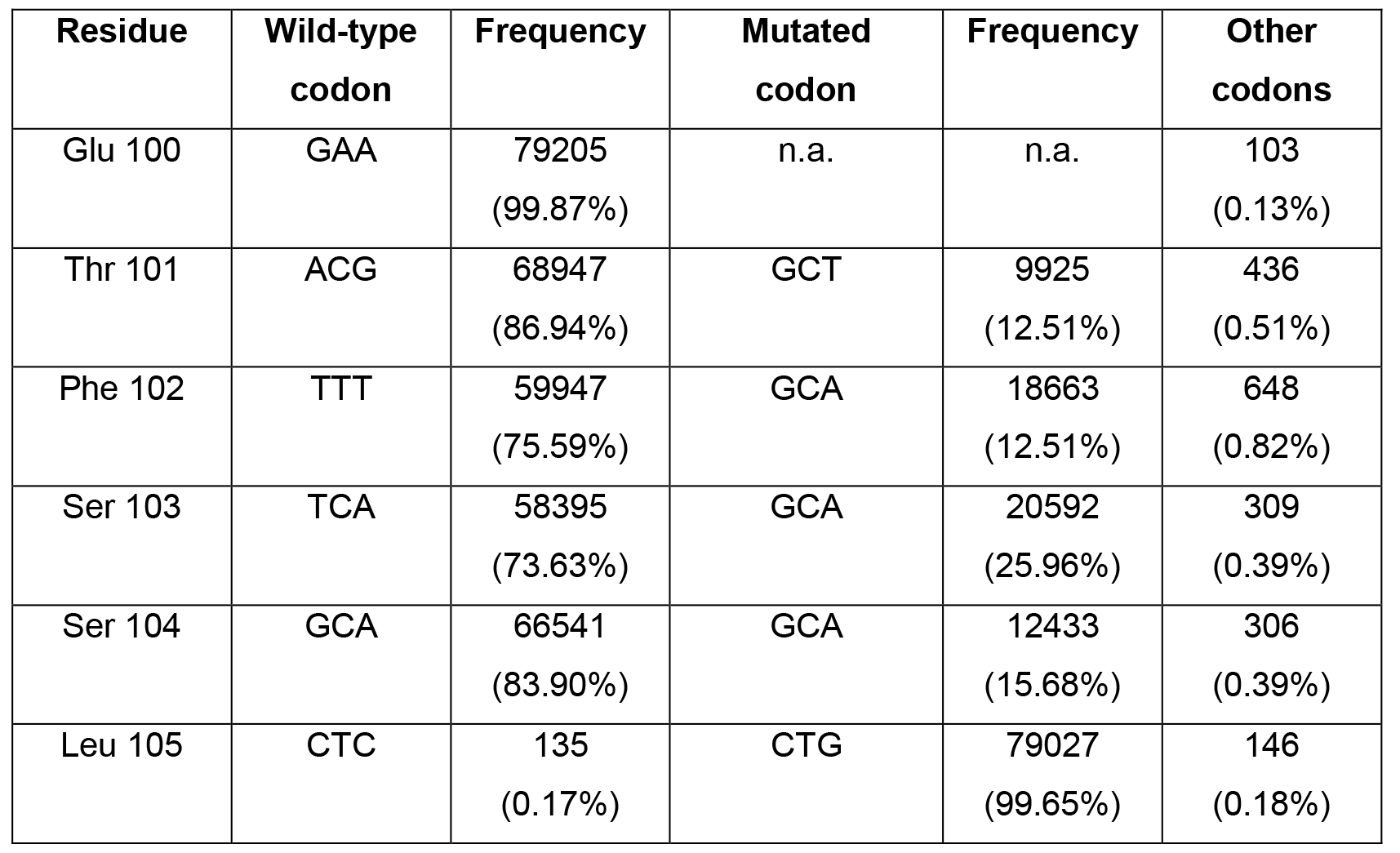
Deep sequencing of CAT gene alanine scanning library. Near complete mutation of Leu105 confirms the efficiency of the approach with alanine represented in all target sites with frequencies between 12.51% and 25.96% (expected frequency: 20%). Sites not targeted by mutagenesis (Glu100 and Trp106) – and where mutations are not applicable (n.a.) – show the underlying error rate of sequencing (0.13 to 0.33%) is comparable with the error obtained in the engineered positions (0.18 to 0.82%).

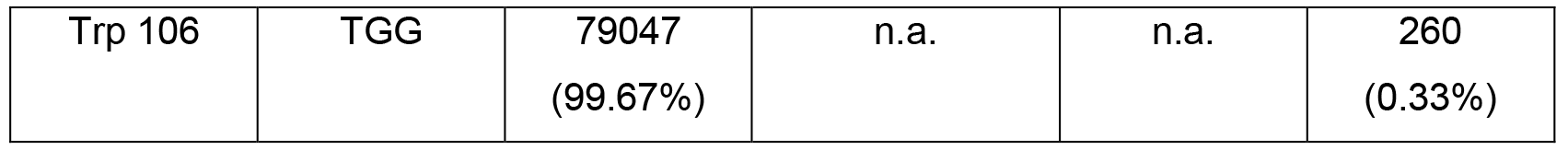

Crucially, an expected proportion of reads (16,739 or 21.1%) were wild-type (containing only the Leu105 recoding). This demonstrates that even though there may be a degree of incorporation bias, with alanine incorporation ranging from Thr101 (12.5%) to Ser103 (26.0%), these are unlikely to have been caused by the mismatches alone.

### Generation of deletion and insertion (indel) libraries

Although insertions and deletions are known to play crucial roles in protein function (e.g. antibodies), there are few tools available to explore length as a parameter in protein engineering(34, 35). We therefore decided to investigate if our Darwin Assembly could be used as a tool to explore indels for protein engineering.

We generated two libraries against the CAT gene: one exploring single and double deletions, and a second one exploring single and double insertions (Figure S2). As with the scanning libraries, the indel libraries were assembled using an equimolar mixture of inner oligonucleotides designed against a common binding site. In addition, similar controls were included, using a wild-type oligonucleotide and the Leu105 (CTC→CTG) controls.

The deletion library was assembled with four inner oligonucleotides, maintaining the wild-type sequence, deleting Phe102 or Ser104, or deleting both sites. The insertion library was generated with three oligonucleotides, maintaining the wild-type sequence, or introducing one or two degenerate codons (NNS) between Phe102 and Ser103.

Assemblies of both libraries were successful and several thousand transformants were obtained for each library. Libraries were pooled and sequenced obtaining approximately 2.1 × 10^5^ reads for analysis. As both libraries contained a wild-type assembly control and were pooled for sequencing, analysis of assembly biases cannot easily be determined.

Nevertheless, there is a clear drop in assembly efficiency between single (16.5% of reads Δ Phe102, 5% ΔSer104) and double deletions (0.39% for ΔPhe102ΔSer104), which we expect reflects the drop in melting temperature of the inner oligonucleotides used in assembly. For the insertion library, it is less clear whether a bias is observed. Importantly, all possible codons encoded by the degenerate inner oligonucleotides were observed for both single and double insertion populations (Table S2-4).

Together, the insertion and deletion libraries confirm that Darwin Assembly can be used to investigate small changes in length in a target gene, but may require further optimisation to minimise assembly biases of deletion libraries.

### Darwin Assembly enables the synthesis of custom complex libraries ideal for directed evolution

Having demonstrated the potential of Darwin Assembly on the CAT gene, we implemented assembly on longer targets: T7 RNA polymerase (2.7 kb) and TgoT DNA polymerase (2.4 kb). These are more typical of enzymes targeted for engineering and represented a greater PCR amplification challenge.

As expected, assemblies longer than 1.2 kb were initially less robust and invariably required considerable PCR optimisation (data not shown). We concluded the PCR problems were caused by carryover of unincorporated inner oligonucleotides and/or partial assembly products from the assembly reaction to PCR. Therefore, we investigated two strategies to remove template and unextended (or partially assembled) oligonucleotides: either purifying assembled molecules (Figure 1) or by selectively degrading template and unassembled products (Figure 2). Both approaches were successful and enabled assembly of libraries spanning more than 2 kb with T7RSS and TgoT respectively.

For T7RSS, we designed a library targeting residues previously implicated in switching promoter specificity(9): Arg96, Lys98, Glu207, Glu222, Asn748, Pro759. The 2724 bp assembly, targeting six codons, used five inner oligonucleotides introducing degeneracy (NNS) at each of the target positions (Figure 4a). Assembled libraries were amplified, cloned and transformed, with approximately 1 × 10^7^ transformants isolated.

Sequencing of the pooled transformants confirmed that at each position, all possible variants were introduced (Figure 4, Table S5). Wild-type contamination, in codons where that could be detected, was always below 0.24% and comparable to sequencing errors detected in non-targeted positions. Incorporation biases were small and correlated primarily with the number of mismatches between template and target codon. Overall, there were also detectable biases over the whole assembly (Kruskal-Wallis ANOVA on ranked target codons, p = 0.001), but since the template residues were clustered in sequence space (5/6 were 67% purine or above) it is possible that the overall incorporation bias observed is a consequence of the individual mismatch biases, rather than a systematic bias.

**Figure 4.**
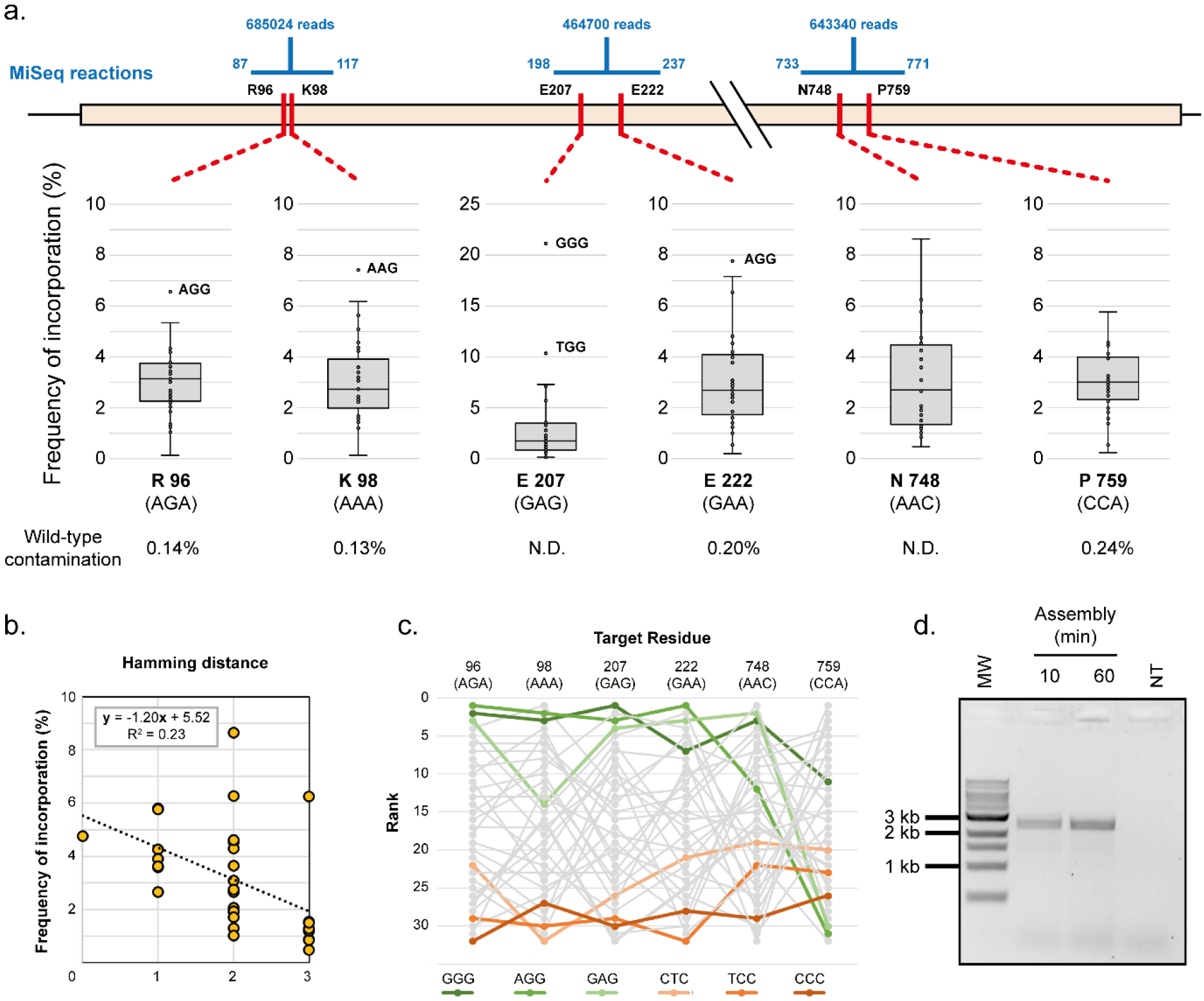
Darwin assembled T7 RNA polymerase library. Sequencing of three separate regions of the assembled gene were used to determine the assembled diversity in the targeted codons (a). Number of reads obtained for each segment are shown as well as the range of each reaction (shown in blue). Frequency of incorporation of the 32 possible codons (NNS) is shown as a box plot, with overrepresented outliers explicitly labelled. Wild-type contamination was determined where possible (N.D.: not determined). Incorporation trends were detected in all positions and best correlated to the number of mismatches they introduced. A typical correlation is shown (b) but other positions varied (0.03 to 0.37), always indicating that the higher the number of mutations being concomitantly introduced (i.e. larger Hamming distance), the lower incorporation of the variant in the population. Incorporation trends were also analysed to determine systematic biases of incorporation. Incorporation frequencies were ranked at each position and rank order analysed (c) – the top three highest ranked (greens) and lowest ranked (oranges) codons are highlighted for clarity. Incorporation biases were detected but it was not clear whether that was the result of the very imbalanced sequence composition of the wild-type residues targeted. Outnest PCR of the T7 RNA polymerase library (expected product of 2840 bp) confirms that as little as 10 minutes of isothermal assembly were sufficient for successful amplification of a library (d). MW: 1 kb ladder (NEB). NT: no template PCR control.

A known limitation of targeting residues with NNS (or NNK) degenerate oligonucleotides is the bias they introduce in translation: oversampling residues that have multiple codons (e.g. Leu and Arg) and increasing the minimum library size required to adequately sample the sequence space introduced. Alternatives to NNS have been developed for site-directed mutagenesis, such as “small intelligent” libraries that mix four oligonucleotides with different degrees of degeneracy (NDT, VMA, ATG and TGG) and in a suitable ratio (12:6:1:1, respectively) to generate equal coverage of all 20 amino acids (36). Such libraries are impossible when targeting multiple sites using simple iPCR. Nevertheless, as Darwin Assembly targets distal sites using independent inner oligonucleotides, it is well suited to generating multi-site intelligent libraries – where nucleic acid variation matches protein variation, maximising library quality.

**Figure 5:**
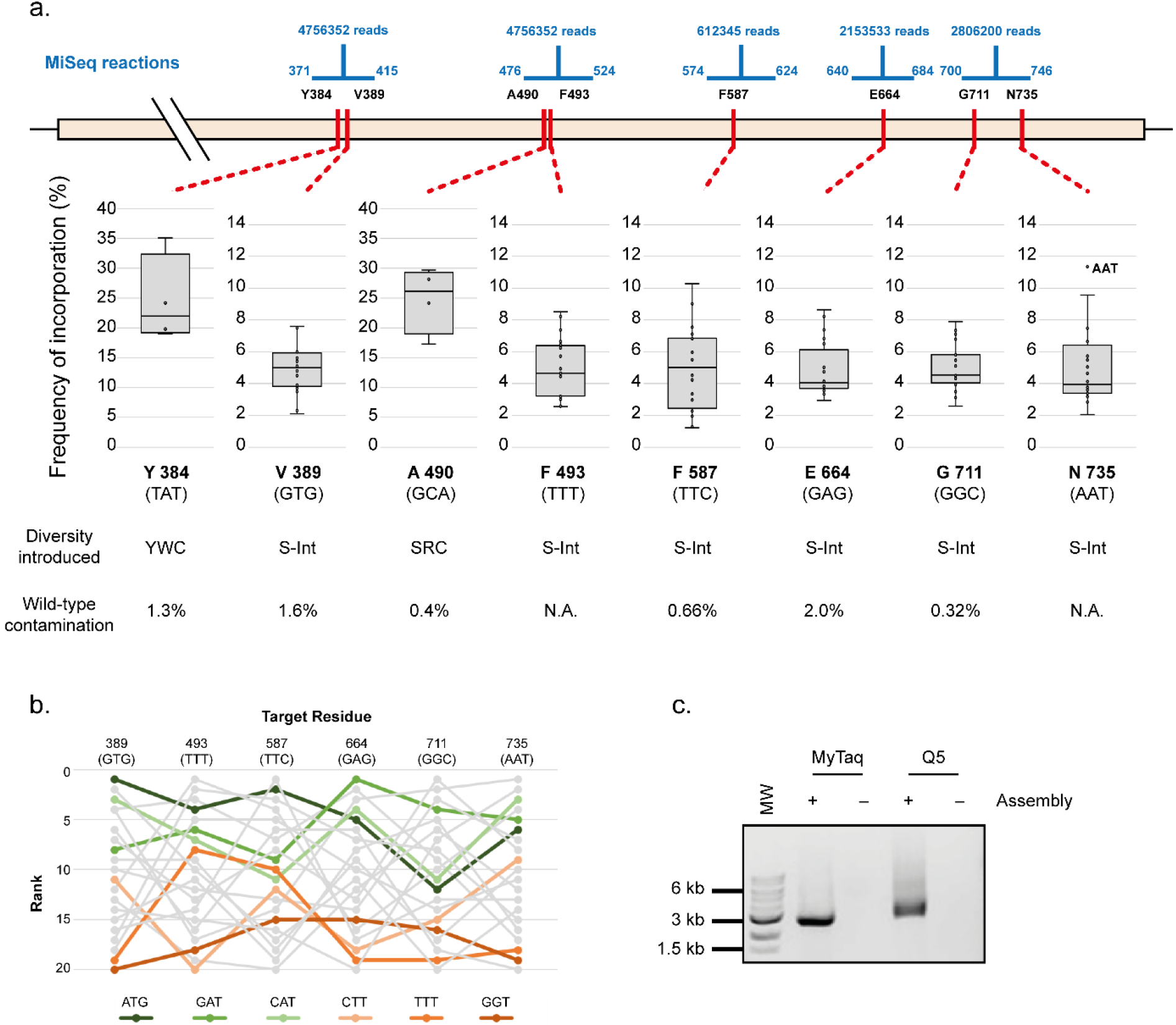
Darwin assembled TgoT DNA polymerase library. (a) Five separate sequencing reactions (range and reads shown in blue) were required to sample the diversity introduced across the eight target residues (shown in red along the TgoT gene). Mutations included focused degeneracies (e.g. YWC used against Y384) or “small intelligent” (S-int) diversity (NDT, VMA, ATG and TGG oligonucleotides mixed in a 12:6:1:1 ratio). Resulting incorporation is shown in box plots with outliers explicitly labelled. Wild-type contamination was determined from positions where diversity excluded those sequences (N.A.: not applicable). As with the T7 RNA polymerase library, incorporation trends and biases were analysed to identify any biases in assembly. Ranked incorporation frequencies are shown for the residues targeted with “small intelligent” diversity, and the top three highest (greens) and lowest (oranges) ranked codons (based on a straight sum of ranks) are highlighted (b). Outnest PCR of the TgoT DNA polymerase library (expected product of 2501 bp) showing that the final PCR can be carried out with either A- (MyTaq) or B-family (Q5) polymerases (c). MW: 1 kb ladder (NEB). NT: no template PCR control.

We therefore designed the TgoT library to target eight residues implicated in template recognition and DNA duplex affinity modulation (Tyr384, Val389, Ala490, Tyr493, Phe587, Glu664, Gly711 and Asn735(10, 37)), with a mixture of limited or with small intelligent degeneracies (Figure 5).

The assembly spanned 2389 bp and targeted the eight residues with six inner oligonucleotides and a non-mutagenic “θ” boundary. The resulting library was successfully amplified, cloned and transformed with an estimated 2.25 × 10^8^ transformants generated. Pooled transformants were sequenced, confirming that the designed diversity was introduced in all targeted residues (Figure 5). As with the T7 library, a positional bias is detectable and correlates weakly to the number of mismatches between template and target codons. Overall, there is a detectable bias in the final assembly beyond a potential systematic error in the mixing of the oligonucleotides (Kruskal-Wallis ANOVA on ranked target NDT codons, p = 0.005), but it is less clear whether the bias is a consequence of the individual template/target codon mismatches – e.g. systematic underrepresentation of GGT even at residue 711 (GGC). (Figure 5b, Table S6-7).

Overall, both T7RSS and TgoT libraries do effectively sample the introduced diversity, given transformants isolated and the measured per site diversity. Both libraries have the expected diversity at multiple distal sites, have minimal contamination of wild-type sequences and are free of assembly errors, despite the size of the genes. Moreover, Darwin Assembly enabled generation of those libraries within a single working day and at a lower cost than current potential commercial suppliers.

## DISCUSSION

Library quality is a major contributing factor to directed evolution(38). It impacts the minimum library size that has to be generated to adequately sample a given sequence landscape(16, 39) and it impacts the choice of selection strategy (depending on the relative rarity of a “successful” variant and gain of function over wild-type). Generating high-quality libraries that target multiple distal sites in a gene is challenging, with commercial library synthesis and modular assembly of the library being available only recently.

Darwin Assembly is a highly efficient method for simultaneous mutagenesis of multiple sites. It is capable of introducing point mutations, bespoke diversity, insertions and/or deletions in complete or partial replacement of wild-type. It is fast, requires no specialised strains or chemical steps and libraries are easily generated in a single working day (from plasmid miniprep to transformation) using commercially available enzymes. We also observed very low levels of wild-type sequence contamination (peaking in TgoT E664 at 1.95% but generally below 0.5%). Overall, this makes Darwin Assembly uniquely suited for generating high quality libraries for directed evolution experiments (38, 40).

Darwin Assembly is amenable to automation, as most steps are enzymatic in compatible buffers and can readily be programmed. We cloned our libraries using type IIS (Golden Gate) restriction cloning, but the freedom to design outnests means any cloning strategy can be employed. Mutations are introduced by oligonucleotides and their design has few constraints – a binding site of at least 11 bases at each end of the “inner” oligonucleotides. As a result, mutation type and density is limited primarily by the oligonucleotide synthetic constraints, which could, for instance, include unnatural bases(41, 42). Similarly, while we only target coding regions of single genes, simultaneous mutation of multiple features (e.g. RBS and gene) or multiple genes (e.g. tRNA and aminoacyl tRNA synthetase) on the same plasmid could be targeted with no protocol modifications, providing the boundary oligonucleotides are sited appropriately.

Post assembly cleanup was not necessary for short fragments, making a one-pot assembly feasible. Longer assemblies greatly benefited from the removal of wild-type template and unincorporated oligonucleotides and both methods we present here were successful. Post-assembly amplification was marginally better for T7RSS, suggesting that purification of assembled fragments may be superior to degradation of template and unused oligonucleotides. We demonstrate that assemblies spanning as much as 2.7 kb are possible and we postulate that maximum assembly length would be determined by the success of post-assembly amplification. In principle, using θ boundary oligonucleotides, it may be possible to replace post-assembly PCR with rolling circle amplification, which would enable assemblies spanning regions longer than 30 kb. We have successfully introduced 38 point mutations covering 19 codons in a single reaction and see no reason why this is the limit of the technology.

## ACKNOWLEDGEMENT

The authors thank Jared Ellefson and Andy Ellington (University of Texas) for the kind gift of pRST.11B.AS3.4 (encoding ScWRS) and the T7RSS gene. We also thank John Chaput (University of California, Irvine) for the kind gift of the pGDR11-KOD plasmid harbouring the KOD DNA polymerase gene.

## SUPPLEMENTARY FIGURES

**Figure S1:**
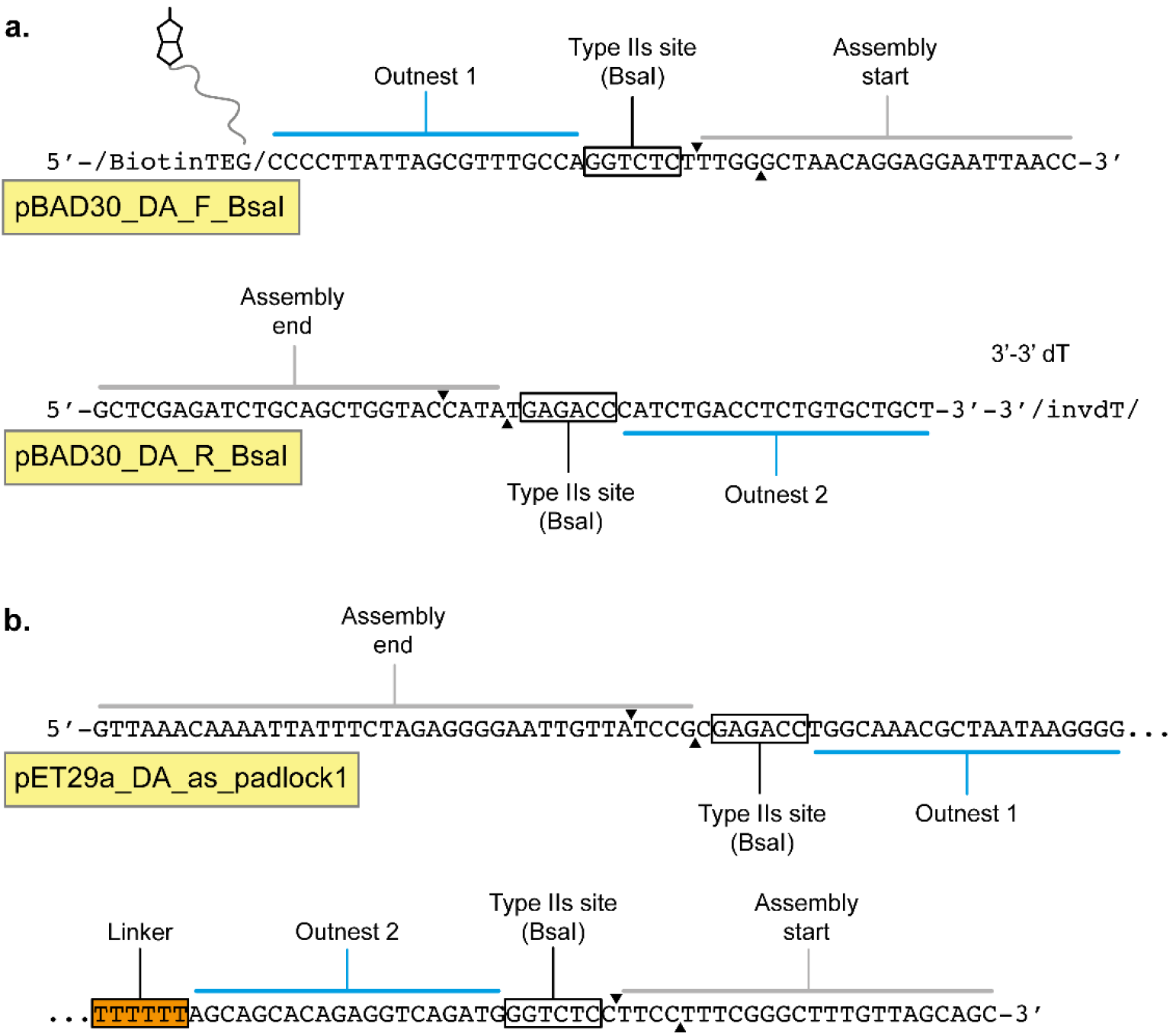
Darwin Assembly boundary oligonucleotide architecture. (a) Design of 2-primer assembly (as described in Figure 1). pBAD30_DA_F_BsaI is the 5’-boundary oligonucleotide used for T7RSS mutagenesis. It is 5’-biotinylated and encodes priming sites (annotated above the sequence) for outnest PCR (blue) and assembly start (grey). The Type IIs recognition site is place between the two priming sites to enable Golden Gate assembly. pBAD30_DA_R_BsaI is the 3’-boundary oligonucleotide used for T7RSS mutagenesis. It encodes the binding site for the assembly end and the reverse complement of Type IIs recognition site and outnest primer used in the downstream amplification. Annotation of reverse complement sequences are shown below the sequence for clarity. This architecture uses the 3’-boundary oligonucleotide to terminate the assembly. Primers shown were used in the assembly of T7RSS libraries. (b) Design of theta assembly oligonucleotide (described in Figure 2). A single oligoucleotide (pET29a_DA_as_padlcok1) was used in the assembly of the TgoT library. Both 5’ and 3’ regions of this oligonucleotide anneal to the plasmid during assembly: the 3’ region as the primer and the 5’ region as the termination point. Between these regions BsaI sites and outnesting priming sites are encoded, linked by a flexible poly(dT) linker (orange), The strand assembled for the TgoT library was the antisense strand and the orientation of the outnest primers was swapped. Oligonucleotide names are shown in yellow boxes and BsaI cut sites are shown as black triangles. Arrows above the sequence represent cuts of the strand being presented, and arrows below the sequence represent the position of the cut sites in the reverse complementary strand generated after the outnest PCR.

**Figure S2:**
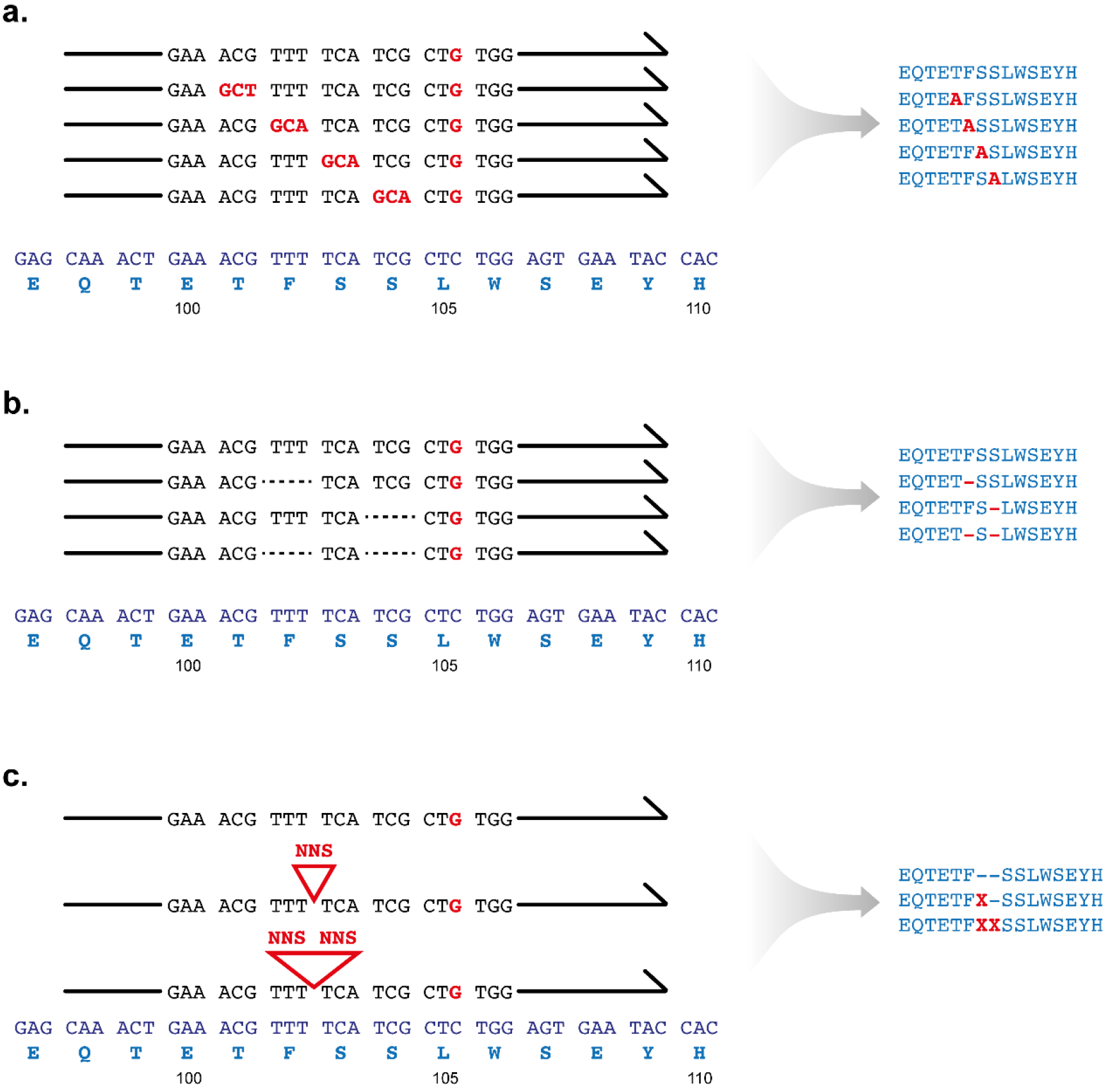
Design of CAT alanine scanning and indel libraries. All oligonucleotides were designed to bind to the same region of CAT gene, with no attempt to adjust overall melting temperature by extending flanking sequences. (a) Five inner oligonucleotides were designed for the synthesis of the alanine scan library, all introducing the control Leu105(CTC→CTG) mutation (and four of them introducing an alanine at one of Thr101, Phe102, Ser103 or Ser104. The oligonucleotides were mixed in equimolar ratio, so all variants were expected at 20%. (b) For the deletion library, four oligonucleotides were designed. All introduced the control Leu105(CTC→CTG) mutation and three of them introduced either single (Phe102 or Ser104) or double (Phe102 and Ser104) mutations. It is likely that the double deletion oligo, resulted in deletion of Phe102 and Ser103 with Ser104 being recoded. (c) For the insertion library, three oligonucleotides were designed introducing Leu105(CTC→CTG) and up to two NNS incorporations between Phe102 and Ser103.

